# Quality assessment of anatomical MRI images from Generative Adversarial Networks: human assessment and image quality metrics

**DOI:** 10.1101/2022.01.03.474792

**Authors:** Matthias S. Treder, Ryan Codrai, Kamen A. Tsvetanov

## Abstract

**Background:** Generative Adversarial Networks (GANs) can synthesize brain images from image or noise input. So far, the gold standard for assessing the quality of the generated images has been human expert ratings. However, due to limitations of human assessment in terms of cost, scalability, and the limited sensitivity of the human eye to more subtle statistical relationships, a more automated approach towards evaluating GANs is required.

**New method:** We investigated to what extent visual quality can be assessed using image quality metrics and we used group analysis and spatial independent components analysis to verify that the GAN reproduces multivariate statistical relationships found in real data. Reference human data was obtained by recruiting neuroimaging experts to assess real Magnetic Resonance (MR) images and images generated by a Wasserstein GAN. Image quality was manipulated by exporting images at different stages of GAN training. *Results*: Experts were sensitive to changes in image quality as evidenced by ratings and reaction times, and the generated images reproduced group effects (age, gender) and spatial correlations moderately well. We also surveyed a number of image quality metrics which consistently failed to fully reproduce human data. While the metrics Structural Similarity Index Measure (SSIM) and Naturalness Image Quality Evaluator (NIQE) showed good overall agreement with human assessment for lower-quality images (i.e. images from early stages of GAN training), only a Deep Quality Assessment (QA) model trained on human ratings was sensitive to the subtle differences between higher-quality images.

**Conclusions:** We recommend a combination of group analyses, spatial correlation analyses, and both distortion metrics (SSIM, NIQE) and perceptual models (Deep QA) for a comprehensive evaluation and comparison of brain images produced by GANs.

## 1. Introduction

Magnetic resonance imaging (MRI) provided a means to understanding the structural and functional heterogeneity of the human brain in health and disease. The recent surge in computing horsepower together with large international collaborative initiatives advanced neuroimaging into a big data science, opening the field to deep learning with a promise for new discoveries [1]. One of the most successful deep learning models has been the Convolutional Neural Network (CNN). Using brain images (e.g. T1-weighted or grey-matter density maps) as input, it has been used in brain age regression [2], brain tumor classification [3], tumor segmentation [4], and a plethora of other applications such as image enhancement, image modality translation, and data augmentation (see [5] for a review).

Many of the latter applications build on generative models which take as input either an image (*image-to-image* approach) or a random vector (*noise-to-image*) and produce an artificially generated brain image as output. The two main architectures for generative models are Generative Adversarial Networks (GANs; [6]) and Variational Autoencoders (VAEs; [7]). GANs have been shown to produce more crisp images than VAEs in diffusion-weighted [8] and T1-weighted images [9] and will be our focus in the rest of the paper. In GANs, the generator creates brain images as outputs using transposed convolution layers, the discriminator is a CNN that tries to classify images as generated or real. Both networks act as adversaries towards each other, with the generator using feedback from the discriminator to create increasingly realistic brain images. In the *image-to-image* approach, the generator receives an input image, typically from a different modality or lower resolution, and produces an output image in a different modality or higher resolution. For instance, GANs have been used to translate T1-weighted images into T2-weighted images [10], Computed Tomography [11], functional MRI [12], and diffusion weighted images [13]. Furthermore, [14, 15] recovered high-resolution images from images reduced in k-space. A potential limitation of the image-to-image approach is that it requires input images from two imaging domains. In the *noise-to-image* approach, realistic MR images are synthesized *de novo*, starting from a random noise vector [16]. Their success rests on the fact that MRIs form a low-dimensional mani-fold and the generator acts as a forward model that maps from this mani-fold to image space. The primary application of noise-to-image GANs has been data augmentation. Several authors successfully used noise-to-image GANs to create realistic T1-weighted 2D image slices [17, 18] or 3D brain images [19, 9, 20]. The noise-to-image approach has also been used in tandem with the image-to-image approach in order to improve tumor detection [21]. *Noise-to-image* problems are arguably harder than *image-to-image* problems, since all anatomical features have to be learned from scratch, including the position and shape of the brain volume and its internal 3D structures.

While much effort has gone into improving generative models, no comprehensive framework for assessing the image quality and biological plausibility of the generated images has emerged yet. The current gold standard for judging visual quality is assessments by neuroimaging experts [8, 22]. In [17] two neuroimaging experts scored images on a 5-points Likert scale with real images scoring higher than generated ones. Several studies used a binary detection task wherein the experts were presented either real vs generated images [22, 8, 18]. Results indicate that generated MRIs are able to mislead experts into believing they are real to a high extent, albeit not perfectly so. Participants reported abnormalities in the shape of landmarks, changes in image contrast, or unusually high symmetry between the two hemispheres as giving away whether or not an image was real.

However, relying on human assessment alone has significant shortcomings. Recruiting human experts is costly, time consuming, and the number of images that can be rated is limited. To avoid this bottleneck and enable fast development cycles, reliable *image quality metrics* are required that can serve as a proxy for human assessment. They can be readily applied to datasets of any size. In image-to-image applications output images can be compared against ground truth reference images using a simple distance metric (see [16] for an overview). A popular metric is L2 loss and quantities derived from it such as mean squared error and peak signal-to-noise ratio (PSNR) have been used for quality assessment [10, 23, 24, 25], as well as Structural Similarity Index Measure (SSIM) [26] and kernel density estimates [22]. A different approach is to use metrics based on intermediate layers of deep learning models. A widely adopted metric in the GAN literature is the Inception Score (IS). However, in [22] IS did not agree well with human perception, since generated images yielded a higher score than real ones.

Additionally, while human assessment and image quality metrics can quantify the extent to which brain images ’look’ realistic, we believe that they provide limited information on their *biological plausibility*. Human observers are sensitive to image artifacts such as checkerboard patterns but other distortions of brain images involve subtle statistical relationships across images that may not be visible to the human eye. For instance, the discovery of structural networks such as the Default Mode Network requires multivariate statistical analyses such as spatial Independent Component Analysis (ICA) performed across dozens or hundreds of MRIs [27]. Furthermore, establishing group differences (e.g. young vs old, male vs female) requires group statistical analysis. It is not clear whether the GAN learns to reproduce such group differences and large-scale structural networks because it is not explicitly trained to do so. We therefore believe that biological plausibility should be investigated as an additional dimension of quality assessment using a combination of group analysis and spatial ICA.

To summarize, the goal of this study was to provide ingredients for a systematic and efficient evaluation of generated brain images. Our contributions are the following:

### 1. Behavioral experiment

We conducted the hitherto largest behavioral study on generated MRIs with 26 neuroimaging experts assessing real and generated grey-matter (GM) density maps from a noise-to-image Wasserstein GAN. Participants performed a detection task (real vs generated images) and a subjective quality rating task using a 5-points Likert scale. To determine the experts’ sensitivity to objective changes in image quality, we exported images at five different stages of GAN training, from early stages (where image quality was supposed to be poor) to later stages. We hypothesized that both detection performance and quality ratings increase with training time and ultimately approach the responses participants give for real images.

### 2. Biological plausibility

For the first time, we performed a detailed analysis of the structural properties of the generated 3D MRIs by performing spatial Independent Component Analysis (ICA) to investigate structural networks and group comparisons that show that GANs are sensitive to gender and age differences. We hypothesized that structural networks and group differences show a high degree of correspondence between real and generated images.

*3. Image quality metrics*. We performed a comprehensive comparison of image quality metrics, ranging from metrics popular in the GAN literature (e.g. Inception Score, Fréchet Inception Distance) to metrics used in image and video quality assessment (e.g. SSIM and VMAF), some of which have not been used in the context of brain images before. Additionally, we trained a Deep Quality Assessment (QA) model on the human data with the goal to provide an automated metric that mimicks human assessment. Since our ultimate goal was to identify a metric that can serve as proxy for human perception, all metrics were tested for their consistency with the behavioral data. We hypothesized that the Deep QA model would outperform distortion metrics such as SSIM and VMAF, especially for high-quality generated images at the end of training.

## 2. Method

The Method section is split into four parts. In Section 2.1, we introduce our generative modeling approach. In Section 2.2, we introduce the behavioral experiment. In Section 2.3 we introduce spatial ICA and group analysis performed in order to investigate the images’ biological plausibility. In Section 2.4, we introduce various image quality metrics and a Deep QA as an objective way of measuring the visual quality of the generated images. For brevity, we refer to any type of brain image derived from an MR sequence (e.g. T1, T2, grey-matter density maps derived from T1) as *MRI* in the rest of the paper.

### 2.1. Generative modeling of MRIs

#### 2.1.1. Data

Models were trained using the Cambridge Centre for Ageing and Neuroscience (Cam-CAN) data [28, 29]. A T1-weighted 3D-structural MRI was acquired with the following parameters: repetition time (TR) = 2,250 ms; echo time (TE) = 2.99 ms; inversion time (TI) = 900 ms; flip angle *α* = 9^°^; field of view (FOV) = 256 × 240 × 192 mm^3^; resolution = 1 mm isotropic; accelerated factor = 2; acquisition time, 4 min and 32 s [29]. The T1 image was initially coregistered to the MNI template, and the T2 image was then coregistered to the T1 image using a rigid-body linear transformation. The coregistered T1 and T2 images were used in a multichannel segmentation to extract probabilistic maps of six tissue classes: gray matter (GM), white matter (WM), cerebrospinal fluid (CSF), bone, soft tissue, and residual noise. The native space GM and WM images were submitted to diffeomorphic registration (DARTEL; [30]) to create group template images. Each template was normalized to the MNI template using a 12-parameter affine transformation. After applying the combined normalization parameters (native to group template and group template to MNI template) to each individual participant’s GM images, the normalized images were smoothed using an 8 mm Gaussian kernel. In total, GM images from 653 participants, aged 18-88 years, were considered as input to noise-to-brain GAN.

It is worth noting that GM maps were chosen over more common modalities such as T1 since our main focus was on the ability of GANs to generate brain structures. In contrast, non-segmented anatomical MR images also involve signals from non-neural structures such as skull, face, and ears. Consequently, it would be impossible to unambiguously attribute human rating and image quality results to brain or non-brain structures. Moreover, spatial brain networks (see Section 2.3) are typically defined using using GM-based morphometry images rather than non-segmented T1 images [31].

Before being fed into the GAN, the following additional preprocessing steps were performed. In the first step, we applied a cuboidal crop on the MRIs thereby removing the empty lateral space that surrounds the brain. Due to the four up-scaling layers in the generator that each up-scale by a factor of 2, the generator was restricted to producing images whose length of any given dimension is a multiple of 2^4^. To bring the size of the real images in line with the generated ones, we resized the cropped MRIs to a final resolution of 96 × 112 × 96. The resizing was necessary in order to fit the MRIs in GPU memory, since the intermittent tensors in the GAN were 5D (input channels x output channels x spatial dimensions) and could consume several gigabytes of video RAM each.

#### 2.1.2. Generative Adversarial Network

In a Generative Adversarial Network (GAN) two neural networks are pitted against each other in a zero-sum game [6]. In this framework, the generator *G* acts as the adversary of the discriminator *D*. Whereas *G* aims to learn the generative distribution of the training data *p*_real_ by approximating it with a generative distribution *p*_gen_, *D* seeks to accurately predict the probability that an image is generated rather than real. The discriminator is a function *D* : ℝ^96×112×96^ → [0, 1], *x* ↦ *D*(*x*) that takes an image as input and returns the probability that this image is real. The generator is a function *G* : ℝ^100^ → ℝ^96×112×96^, *z* ↦ *G*(*z*) that takes a random vector *z* sampled from a probability distribution *p*_*z*_ over ℝ^100^ and returns an MRI. Training is designed to maximize the quality of the generated images by learning the generative distribution *p*_real_. The model training reaches optimality when *D* is unable to discriminate between real and generated samples. The objective of the discriminator is governed by the equation

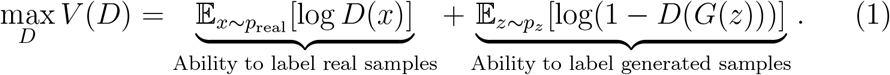

*D* seeks to maximize the probability for samples *x* from the training data and minimize the probability for samples generated from a random vector *z*. This formulation encourages the discriminator to become sensitive to image features that tell the difference between real and generated images. The generator is governed by the equation

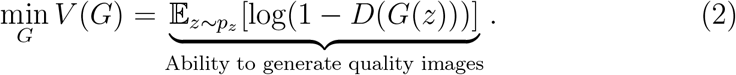

*G* seeks to maximize the probability that its generated images are labeled as ‘real’ by *D*. Both objectives can be combined into a joint loss function that is used for GAN training and optimized using alternating gradient descent

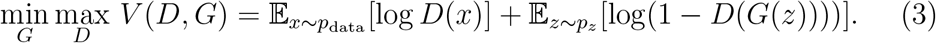

Since *G* learns a mapping from a low dimensional vector space to a high dimensional output, the space from which the input vector originates is a representation of the low-dimensional manifold of MRIs. This quantity is a lower bound of the Jensen-Shannon divergence between the real and generated distributions. GAN training using this objective function can fail to converge due to near-zero gradients when the true and generated distribution do not overlap and also. A more stable alternative that provides useful gradients for non-overlapping distributions is given by Wasserstein GANs (WGANs) [32, 33].To this end, the objective function is modified to the Earth Mover or Wasserstein-1 distance between *p*_real_ and *p*_gen_ given by

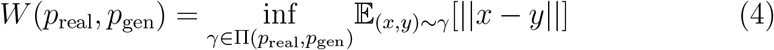

where Π(*p*_real_, *p*_gen_) is the set of joint distributions whose marginals are given by *p*_real_ and *p*_gen_. Practical implementations of this loss function are given in [32, 33].

#### 2.1.3. Generating MR images using GANs

The structure of the noise-to-brain is shown in Figure 1. Note that the output of each convolutional layer is 4D (3 spatial dimensions and 1 output channels dimension). For displaying purposes, it is shown as 3D in the figure.

**Figure 1:**
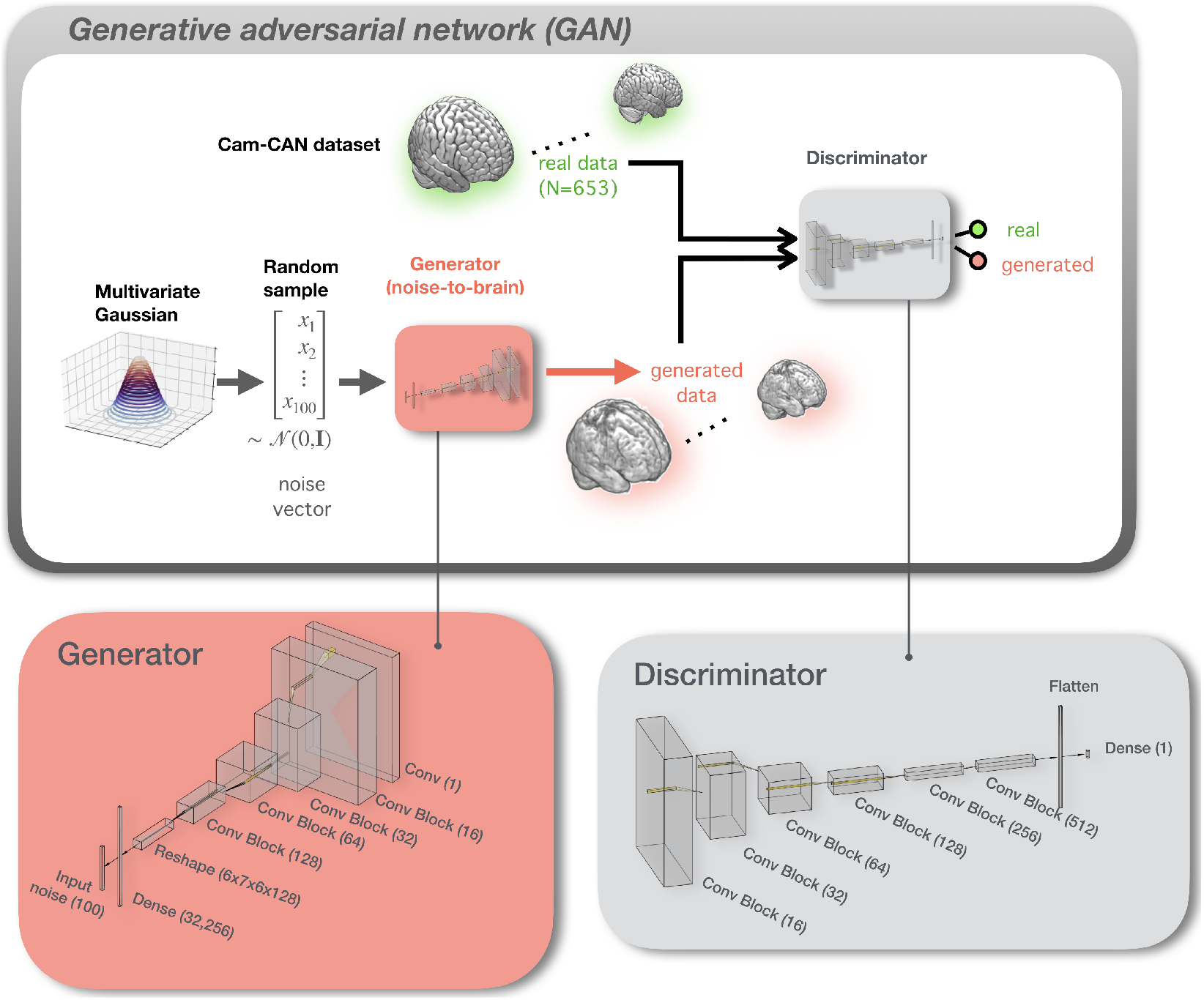
Structure of the noise-to-brain GAN. For the generator and discriminator, the number of filters is given in brackets, whereas for a dense layer, the number of units is shown in brackets.

### 2.2. Behavioral experiment

#### 2.2.1. Participants

Forty-five participants volunteered in the online study. Out of these, 26 participants (10 female, 16 male) completed all parts of the study (questionnaire, detection task, and subjective rating task). Only complete datasets were used for further analysis. Participants were aged 18 to 62 years (*µ* = 33.12, *σ* = 8.48), with the majority of participants (16) in their 30s. All participants were neuroimaging experts (e.g. neuroscientists, neurologists, radiologists) at various levels of seniority (from undergraduate student to professor). They did not receive remuneration for their participation. Due to COVID-19 related restrictions (most of the data was collected during UK lockdowns) and the difficulty of recruiting a sufficient number of MR experts locally, the study was conducted online. Participants were mainly recruited through the authors’ research networks, although no contact details were recorded. Participants gave written informed consent using an online form. Ethical approval was obtained from the COMSC Research Ethics Group at Cardiff University (COMSC/Ethics/2020/034).

#### 2.2.2. Behavioral experiment

As stimuli, we used middle slides in the sagittal, coronal, and transverse plane extracted from the 3D images. These slices were horizontally concatenated to 432 × 288 pixels images. An image was either based on the real MRI data or generated by the GAN. To sample the generated images at different stages of GAN training, ’fake’ images were drawn from five different batch numbers: 344, 1055, 7954, 24440, and 60000 (final batch).

The experiment was conducted online using PsyToolkit [34, 35] (see Figure S1, Supplementary Material, for a visual depiction). Participants carried it out on their own computer by following a weblink. After filling out a brief questionnaire to indicate their expertise and familiarity with MRI data, the experiment started in fullscreen mode. It took about 15 minutes and was split into three phases: In the training phase, participants performed a practice detection task to familiarize themselves with the stimuli. The next two phases were the detection task and the subjective rating task. The order of these two phases was randomized across participants. Each phase was preceded by a 3 s countdown.

Training consisted of 30 randomized trials. In each trial, a real or generated (’fake’) MR image was presented at the center of the screen. Participants were instructed to quickly and accurately indicate their choice with the arrow keys on their keyboard, using the left index finger for the left arrow key (‘real’) and the middle finger for the right arrow key (‘fake’). The presentation time of the image was unlimited, although for practical reasons the trial ended after 20 s and it was marked as timed out. After the trial, feedback was presented (’correct’, ’wrong’, ’time out’). Trials were separated by an inter-trial interval of 400-600 ms. Button press and reaction time were recorded. The proportion of stimuli was 1/3 real (10 images) and 2/3 fake (20 images) with equal proportions for each of the five ’fake’ levels (4 images each).

The detection task was structured just like the training task but no feedback was presented. There were 240 trials split into three blocks of 80 trials. The proportion of stimuli was 1/3 real (80 images) and 2/3 ’fake’ (160 images) with equal proportions for each of the five ’fake’ levels (32 images each). Participants were asked to take a break after every block. The experiment resumed upon key press.

The subjective rating task consisted of 30 trials comprising 5 real images plus 5 images per batch. Participants were rated their visual quality by selecting an item from a 5-points Likert scale with the options “very real”, “relatively real”, “neutral”, “relatively fake”, and “very fake”. The options were presented below the image and chosen via a mouse click. Presentation time was unlimited although for practical reasons the trial ended after at most 10 s. Across the experiment, all images were unique. Once it was finished, participants had the option to submit a comment in a text box.

#### 2.2.3. Behavioral data processing

For the detection task, the proportion of ’real’ responses, proportion of correct responses, and reaction time (RT) were investigated. Trials with timeouts or *<*150 ms RT were discarded. One participant was removed due to a large amount of timed out trials (*>*10%). For every participant, mean ’real’ responses and correct responses were calculated by averaging across all images corresponding to a given batch. For the RT analysis, mean RTs were calculated using a trimmed mean approach wherein the 5% trials with the largest RTs were excluded and mean RT was calculated across the remaining trials [36]. For the subjective rating task, Mean Opinion Scores (MOS) were calculated by averaging across all images in a batch. RTs were processed in the same way as for the detection task.

### 2.3. Group analysis and spatial Independent Components Analysis

To test whether the GAN generates biologically plausible data we compared the similarity between generated and real data using data-driven and model-based analyses. For the model-based approach, we sought to reproduce subtle differences between groups wherein we trained separate GANs on different subsets of the data. Training separate models for different groups is an approach that has been explored before [37]. To investigate age effects, we split the data into elderly (*>* 70 years; 172 MRIs) and young 287 (*<* 40 years, 185 MRIs). To test for effects of sex, we contrasted male (323 MRIs) and female (330 MRIs) MRIs. Once trained, we randomly generated as many MRIs as were in each of the subsets. The hypothesis was that if the GAN is sensitive to group characteristics, the generated MRIs should display similar age and gender effects as the real MRIs. For the data-driven approach, we used independent component analysis to decompose the synthetic data in a small set of spatially independent maps that correspond sensibly with known neurobiological relationships in the real data [31]. In particular, spatial ICA was implemented on real and generated GM maps separately. For each dataset, data were decomposed to a small number of spatially independent sources using the Source-Based Morphometry toolbox

[38] in the Group ICA for fMRI Toolbox (GIFT)^1^. By combining the PCA and ICA, one can decompose a participants-by-voxels into a source matrix that maps independent components (ICs) to voxels (here referred to as “IC maps”), and a mixing matrix that maps ICs to participants. The voxels that carried similar information across participants would have higher values and group to a set of regions. The spatial IC maps were then converted to z-scores. The correspondence between real and generated MRIs was based on the spatial correlation between thresholded IC maps, where the threshold was set to z-value *>* 3. To further explore that the results were independent of the selected number of components, we repeated the procedure for a range of different components (5, 10, 15, 20, 25, 30).

### 2.4. Image Quality Metrics and Models

#### 2.4.1. Image quality metrics

In pursuit of an automated metric that can serve as a proxy for human assessment, we surveyed a range of image metrics. For the sake of comparability we applied these metrics to the same 2D images that were used in the behavioral experiment.

We first calculated four metrics widely adopted in the GAN literature, namely Inception Score (IS), Modified Inception Score (MIS), Fréchet Inception Distance (FID), Maximum Mean Discrepancy (MMD) [39]. In addition, we used an array of metrics developed by the image/video processing community that are supposed to approximate human quality assessments [26]. We tested six among the most widely adopted metrics in the image quality assessment field, Peak signal to noise ratio (PSNR), Structural Similarity Index (SSIM), Structural Similarity Index (MS-SSIM), Video Multimethod Assessment Fusion (VMAF), Natural Image Quality Evaluator (NIQE), Blind/referenceless image spatial quality evaluator (BRISQUE); see [40] for a review). In the following, each of these metrics is introduced in more detail. As before, *p*_real_ and *p*_gen_ are used to denote the distributions of the real and generated images. Furthermore, *p*(*y*) denotes the distribution of the class labels and *p*(*x*|*y*) the distribution of images in class *y*.

- *IS* is a popular GAN metric using a CNN pre-trained on ImageNet [41]. We used the InceptionV3 model trained with 1000 classes. Its final layer is a softmax layer that represents the conditional distribution *p*(*y* | *x*) of class labels *y* given an input image *x*. IS is then given by the formula

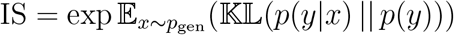

where 𝔼 is the expectation, *x ∼ p*_gen_ are the images sampled from the generator, 𝕂𝕃 refers to the Kullback-Leibler divergence, and *p*(*y*) is the marginal distribution of class labels. IS favours generated images that show a clear class membership, characterized by conditional class labels *p*(*y* | *x*) with a ‘peaky’ (low entropy) distribution, at the same time favouring a broad coverage of multiple classes characterized by a ‘flat’ (high entropy) marginal distribution *p*(*y*). A detailed discussion of IS is provided in [42].
- *MIS*, unlike IS, takes into account the desire for diversity of images within a given class. To achieve this, [43] incorporated a cross-entropy term −*p*(*y* | *x*_*i*_) log *p*(*y* | *x*_*j*_) into IS that represents within-class diversity yielding the modified Inception Score

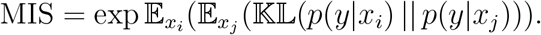
- *FID* uses the feature embeddings of a CNN wherein it models the distributions of real and generated images as multivariate Gaussians [44]. FID is then given as the Fréchet distance between these two Gaussians,

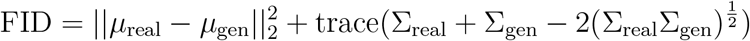

where *µ*_real_ and *µ*_gen_ are the means for real and generated images, and Σ_real_ and Σ_gen_ are their respective covariance matrices. We calculated FID using InceptionV3’s penultimate layer. The activation maps were condensed into 2,048 features by applying global average pooling.
- *MMD* measures the distance between two distributions, operationalized as the distance between its mean embeddings in feature space. Gretton et al. [45] implemented MMD using kernels, yielding

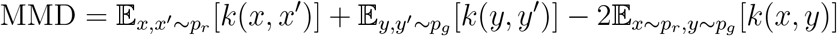

where *k* is a kernel function. We calculated MMD by again using the penultimate layer of InceptionV3 and the Radial Basis Function (RBF) kernel 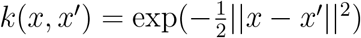.
- *PSNR* is a simple metrics based on the pixel-wise difference between a target image *x* and a reference image *y*,

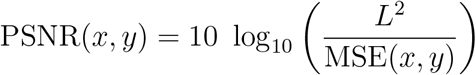

where MSE is the mean-squared error between target and reference and *L*^2^ is the maximum pixel value (e.g. 1.0 for data normalized in the 0-1 range).
- *SSIM* quantifies similarity using a product of three separate terms for luminance, contrast, and structure similarity [26]. The product can be represented as

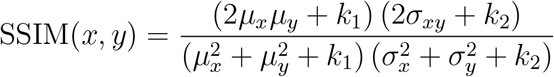

where *µ*_*x*_, *µ*_*y*_ are the mean pixel values of the target and reference images, *σ*_*x*_, *σ*_*y*_ are their variances and *σ*_*xy*_ is the covariance, and *k*_1_, *k*_2_ are regularization hyperparameters used for numerical stability. We computed SSIM using TensorFlow’s implementation with default hyperparameters (11×11 Gaussian filter with width 1.5, *k*_1_ = 0.1, *k*_2_ = 0.3).
- *Multi-scale Structural Similarity Index (MS-SSIM)* is an extension of SSIM that aims towards scale invariance [46]. The image is analyzed at its original scale and at multiple low-pass filtered and downscaled versions of it. The SSIM values from the different scales are then combined into a single estimate. We used the VMAF command line tool to calculate MS-SSIM.
- *VMAF* was originally developed for video quality assessment [47]. The VMAF model has been trained on a Netflix quality assessment dataset wherein viewers scored videos on a 5-points Likert scale summarized by Mean Opinion Scores (MOS) and normalized to the 0-100 range. We used the VMAF command line tool (github.com/Netflix/vmaf). Images were padded to 480 × 360 px and then exported to yuv format using FFmpeg (www.ffmpeg.org).
- *NIQE* computes image statistics based on normalized luminance values. A multivariate Gaussian distribution is fit to the image statistics and its Mahalanobis distance to a fit obtained on a corpus of natural images is calculated

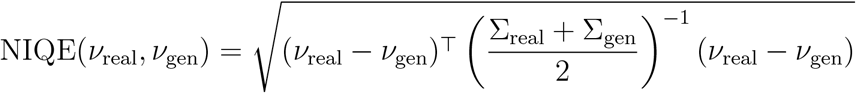

where *ν*_real_ and Σ_real_ are the mean feature vector and covariance matrix obtained on the corpus and *ν*_gen_ and Σ_gen_ are the corresponding quantities computed on the image that is being assessed. We also used a second version of NIQE referred to as NIQE-MRI wherein the model was pretrained on an independent set of MRIs. Matlab’s niqe and fitniqe functions were used.
- *BRISQUE* is a metric that, in contrast to NIQE, requires reference images with distortions as well as MOS to train a SVR model on the image statistics [48]. Just as NIQE, we also pretrained a second version of the model on MRIs. Since BRISQUE requires MOS it was trained on the subjective rating data and cross-vaidation was used to obtain unbiased scores for the images. Matlab’s brisque and fitbrisque functions were used.

Note that IS, MIS, NIQE and BRISQUE are reference free metrics. Therefore, we could calculate scores for both generated and real images. For the other metrics, we used generated images from each of the batches as target images and the real MRIs as reference images. FID and MMD only require a distribution of generated images. However, PSNR, SSIM, MS-SSIM, and VMAF are full reference metrics, that is, for each generated image they require a matched real image. Since there was no matching between generated and real images, we considered all possible pairs of real/generated images. Another distinction between the metrics is that IS, MIS, PSNR, SSIM, MS-SSIM, and VMAF are quality metrics (better quality = higher value) whereas FID, MMD, NIQE and BRISQUE are distance metrics (better quality = lower value).

#### 2.4.2. Deep QA model

An alternative to image-based metrics is to train a model that tries to mimick human perceptual assessments. Such perceptual models have been used both for MRIs and in the wider image/video quality literature [49, 50]. We implemented two CNNs that essentially performed the same task as the human experts, namely classifying images as ’real’ or ’fake’ (detection task) and assigning a subjective rating to an image on a 5-points Likert scale (rating task). The model architecture is depicted in Figure 6a.

##### Detection task model

The first part of the model consisted of a InceptionV3 model pretrained on ImageNet with the classification layer removed. The InceptionV3 output was fed into two dense layers (32 and 16 units, dropout rate 0.1, leaky ReLU activation with *α* = 0.1), denoted as MRI features. The final layer was a single sigmoid unit. The model was trained on a set of 3611 real and generated images not used in the behavioral experiment. Images from all batches were pooled and labeled as ’fake’. The model was initially trained for 5 epochs with the InceptionV3 model frozen. Subsequently, the last two convolutional layers of InceptionV3 were unfrozen and training continued for another 20 epochs. Binary cross-entropy was used as loss function with an Adam optimizer, a learning rate of 10^−4^ (reduced to 10^−5^ after 5 epochs), and a batch size of 16. Finally, the model was tested on the 240 images used in the detection task and the predicted logits were recorded for each image. No behavioral data was used.

##### Rating task model

The challenge of training a model on the rating task was the small number of only 30 images. Therefore, we explored a transfer learning approach wherein we first trained the detection task model and then fine-tuned it on the rating task data. To this end, we took the detection task model and replaced the sigmoid by a softmax layer with 5 units representing the Likert scale ratings. The 30 rating task images were split into train and test sets using 5-fold cross-validation. For the training images, class labels (representing ratings) could not be assigned unequivocally since there were disagreements between the human raters. To encode this uncertainty in the model we used probabilistic class labels: For each image, the empirical distribution of ratings was determined across raters and used to initialize a probability distribution. In each batch, class labels were randomly sampled from these distributions. The model was initially trained for 20 epochs with the InceptionV3 model and the MRI features frozen. After this, the MRI features were unfrozen and model was trained for another 200 epochs. Categorical cross-entropy was used as loss function with an Adam optimizer, a learning rate of 10^−4^ (reduced to 10^−5^ after 20 epochs), and batch size 6. Both models were trained 100 times with randomly initialized weights. Results were averaged across runs.

#### 2.4.3. Hardware and software

GANs were trained on NVIDIA Tesla P100 GPUs with 16 GB VRAM using Supercomputing Wales with TensorFlow 1. Statistical analyses were performed on a standard Desktop computer using Python 3.6 with the packages Statsmodels 0.13, Scipy 1.5.2, and TensorFlow 2.4, as well as Matlab R2018b.

## 3. Results

### 3.1. Qualitative analysis of generated MRIs

Figure 2 shows a 3D rendering of the generated images using MRIcroGL [51]. To indicate how the images evolve across the different stages of training, Figure 2a shows images for six logarithmically spaced batch numbers ranging from an early batch (batch 11) to the final batch (batch 60000). Each of the six renderings is based on the same random vector. The GAN initially outputs a block of noise (batches 11 and 90) from which a brain is eventually carved out (batches 344 and 843). After this, GAN training seems to focus on improving details and removing noise voxels outside the brain volume (compare batches 3,240 and 60000). More incidental evidence for the GAN devoting a lot of computational resources to cancelling noise outside the brain is provided in Figure S2 (Supplementary Material). Figure 2b shows an ‘exploded’ version of a generated MRI followed by two different random examples with a 3D view and various planar views. The model is able to convincingly mimic the overall shape of the brain and prominent structures such as lateral fissure and the cerebellum. The ‘exploded’ view indicates that the internal grey-matter structure is reproduced as well. The shortcomings of the GAN become more evident when viewed side by side with a real MRI example in Figure 2c. Here, large-scale structures such as the lateral fissure are more regular and the image looks less noisy overall.

**Figure 2:**
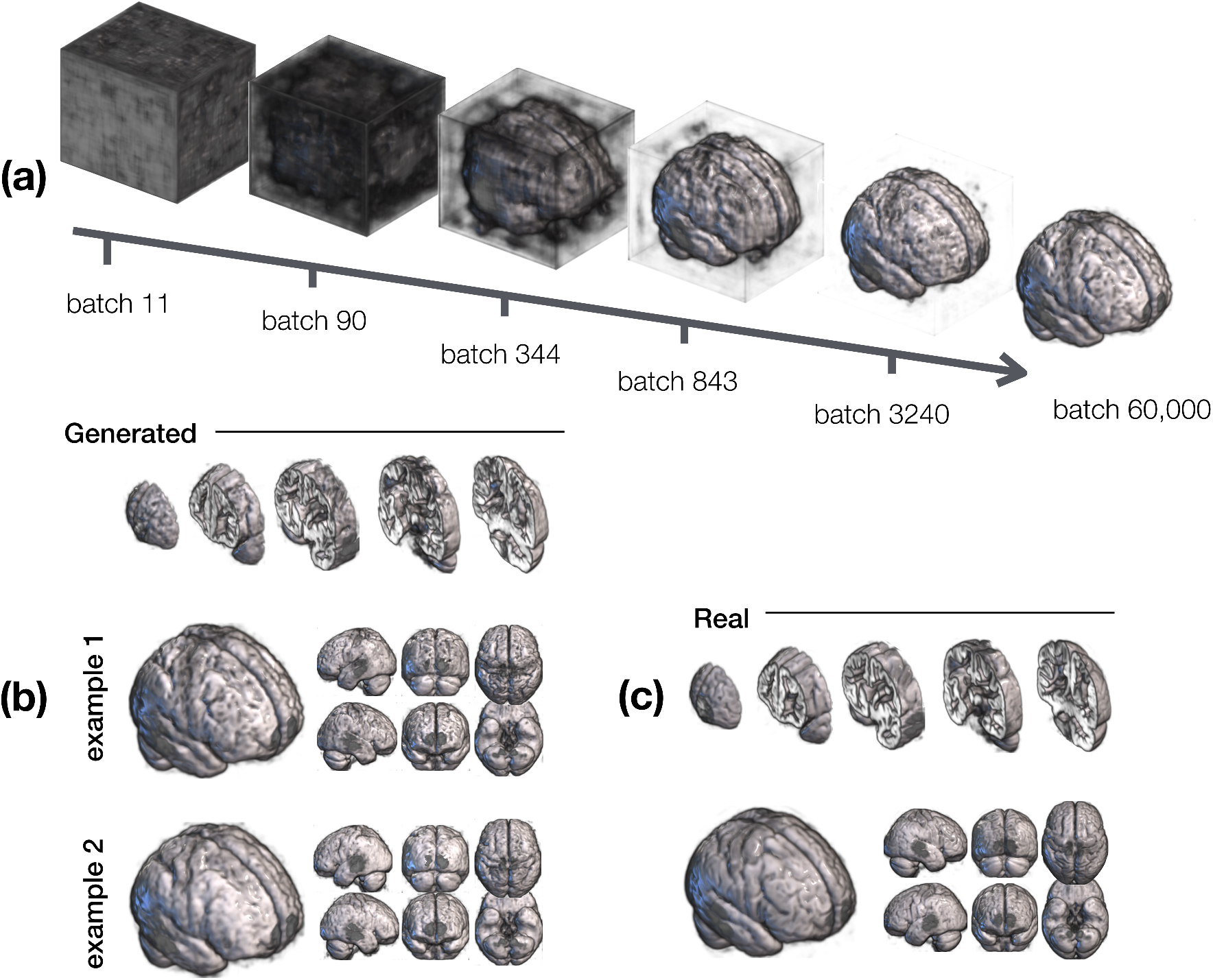
Qualitative results showing a MRIcroGL rendering of the generated 3D images. (a) Generated images at six different times (batch numbers) during GAN training. Whereas only a volume of noise is generated in the initial stages (batches 11 and 90), the basic brain structure is evident from batch 843. Much of the subsequent training appears devoted to refining details and removing the noise outside the brain. Batch 60000 corresponds to the final model. (b) ’Exploded’ view of a generated MRI in the first row indicates that the GAN correctly reproduces the internal structure of the brain, followed by two examples for generated MRIs. (c) For comparative purposes, the same rendering for a real MRI is shown. Comparing real and generated MRI, we observe that the generated image looks slightly more noisy and irregular, especially for large elongated structures such as the lateral fissure.

### 3.2. Correlation analysis and spatial ICA

To ground these observations in a more quantitative analysis, we performed correlation analyses, an analysis of age and sex group effects, and a spatial ICA analysis, depicted in Figure 3. Correlations were calculated across all brain voxels between pairs of MRIs. To account for possible effects of small spatial misalignments and high-frequency noise, correlation analyses were repeated for images post-processed with a 2D Gaussian smoothing filter (*σ* = 3 voxels).

**Figure 3:**
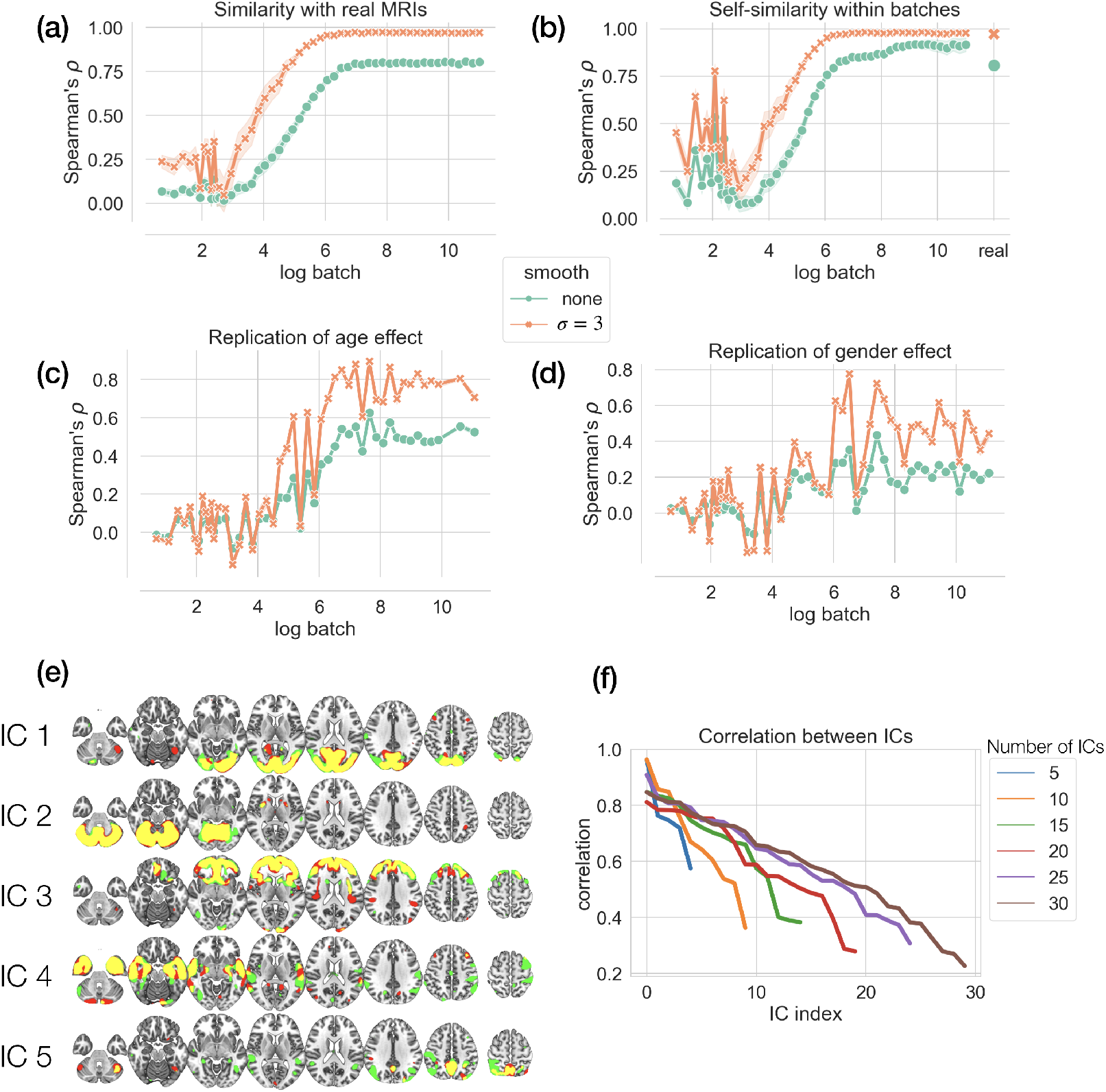
Correlation analyses and spatial ICA. (a) Similarity between generated and real 3D MRIs measured as their average correlation across voxels as a function of log batch number. The shaded area corresponds to 1 standard deviation. Results shown for original data without smoothing (green) and smoothed data (Gaussian kernel, *σ* = 3, orange). (b) Within-batch similarity, average correlation between pairs of images within a batch. Correlation within real MRIs is shown as an additional data point on the right. (c) Replication of the age effect (’old minus young’). (d) Replication of sex effect (’male minus female’). Correlation starts around zero and increases significantly with training, settling at a significant but moderately low value of 0.17. Smoothing significantly boosts correlation (0.42). (e) Spatial ICA maps for ICA with 5 components. Components for both real and generated MRIs have been matched and overlayed. Color coding indicates whether a voxel belongs the real (red) or generated (green) MRIs, with the intersection shown in yellow. The large overlap suggests significant spatial correspondence between real and generated MRIs. (d) Correlations between real and generated ICs for ICAs with different numbers of components (between 5 and 30). High degree of correspondence for the first few components is consistently found.

Figure 3a depicts Spearman’s rank correlation *ρ* between generated and real MRIs. Every generated image was paired and correlated with every real image, then mean and standard deviation were calculated across all pairs. Correlation increased significantly then saturated relatively early in training. Since we masked out non-brain voxels, this early saturation nicely dovetails with our earlier observation that the late stages of GAN training seem largely devoted to removing noise outside the brain. The correlation for the final model was 0.8. Smoothing with a Gaussian kernel with a standard deviation of *σ* = 3 voxels yielded a correlation of 0.97, a statistically significant increase (Wilcoxon Rank Sum test, *z* = 799.76, *p <* 0.0001). The significant effect of smoothing suggests a high degree of correspondence between the large-scale structure of generated and real MRIs.

Figure 3b depicts the correlation among MRIs within batches. Correlation among real MRIs has been added as additional data points (separate rightmost data points). Correlation started low and quickly saturated at a high level. For the final model the correlation amounted to 0.92, whereas it was 0.98 with smoothing, a significant increase (*z* = 563.47, *p <* 0.0001). Corresponding correlations for real MRIs were 0.81 without and 0.97 with smoothing, again a significant difference (*z* = 565.08, *p <* 0.0001). Directly comparing generated and real MRIs, correlation was significantly higher for generated MRIs both for unsmoothed (*z* = 549.74, *p <* 0.0001) and smoothed (*z* = 350.98, *p <* 0.0001) data.

Figure 3c depicts how well the GAN replicates the age effect found in real data. In real data, the age effect was operationalized as the average MRI in the elderly group (*>* 70 years) minus the average MRI in the young group (*<* 40 years). To generate MRIs specifically from these groups, two different GANs were trained, one on the elderly MRIs and one on the young ones. Since the age effect was a group effect, correlations for individual MRIs were not available. Therefore, to calculate errorbars, 100 different bootstrap samples of the GAN data were created. Correlation between real age effects and the generated age effects increased with batch number, suggesting that the GAN was sensitive to these group characteristics. In the final batch, correlation was 0.52 for the unsmoothed and 0.71 for the smoothed data, a significant difference (*z* = 12.22, *p <* 0.0001).

Figure 3d depicts the replication of a gender effect, operationalized as the average ’male MRI’ minus the average ’female MRI’. Again, two different GANs were trained on male and female MRIs separately and bootstrapping was used to calculate errorbars. Correlation increased with batch number, again suggesting that the GAN is sensitive to these group characteristics. However, it settled at a lower value than for the age effect. In the final batch, correlation was 0.22 for the unsmoothed and 0.44 for the smoothed data, a significant difference (*z* = 12.22, *p <* 0.0001).

Figure 3e depicts 5 ICs for a spatial ICA with 5 components. ICs in real and generated MRIs were matched by their correlation. To binarize the maps, one-sample t-values were calculated for all participants and thresholded at a t-value of 3. A large amount of overlap (shown in yellow) is evident especially for the first few ICs. Non-overlapping voxels are shown in red for real and green for generated data. Figure 3f depicts the correlations between IC maps for different spatial ICAs using a different number of components (5, 10, 15, 20, 25, 30). For all analyses, we found correlation values *>* 0.57 for the first five components, suggesting a moderate to high degree of consistency of components for real and generated MRIs.

### 3.3. Behavioral results

Figure 4 depicts the behavioral results. For both tasks, reaction times (RTs) increased with batch number, approaching the RT for real images (Figure 4a and d). For the detection task, the percentage of ‘real’ responses increased with the batch number, although it stayed short of the corresponding number for real MRIs (Figure 4b). For completeness, the correct score is also depicted in Figure 4c. For the subjective rating task, Mean Opinion Scores (MOS) are shown, that is, averages when ratings are encoded as integer numbers 1 through 5. MOS increased with batch number but did not reach the MOS obtained for real MRIs.

**Figure 4:**
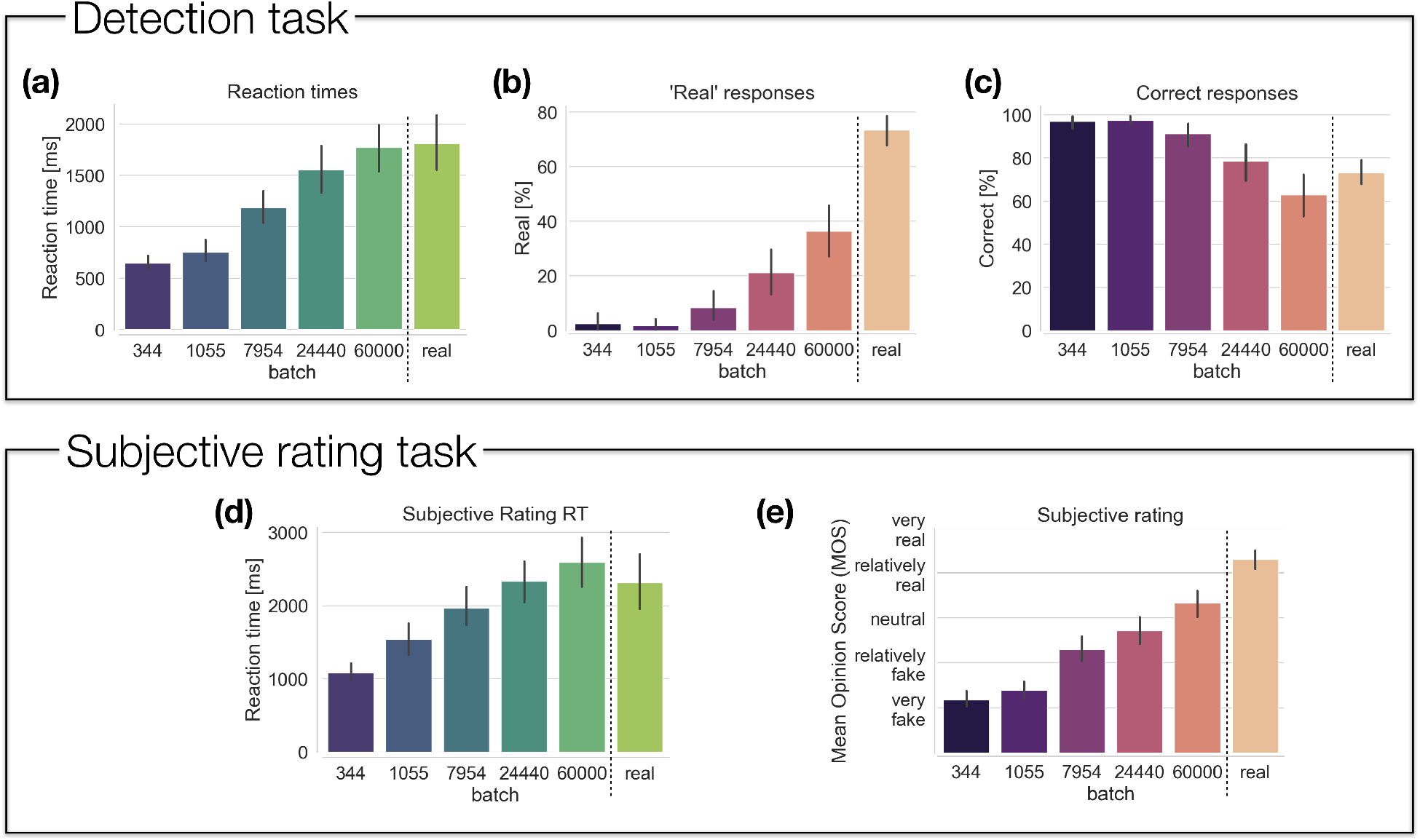
Behavioral experiment results for the detection task and subjective rating task. Results are shown for generated MRIs from different training batches (indicated by the batch numbers) and for real MRI data. (a) In the detection task, reaction times (RTs) increased with batch number, approaching the RT obtained for real MRIs. (b) Percentage ‘real’ responses increased with batch number but stayed short of the proportion obtained for real data. (c) Percentage correct responses decreased with batch number, indicating that it becomes harder to distinguish real and generated MRIs. (d) In the subjective rating task, RTs increased with batch number. For batch 60000 RT slightly overshot the RT found for real MRI. (e) Subjective ratings, here summarized as Mean Opinion Scores, increased with batch number albeit staying below the rating for real MRIs for batch 60000.

To substantiate these observations statistically, we used linear mixed-effects models (LMMs) with a fixed effect (log batch number) and a random effect (participant). LMMs are useful in the case of correlated responses arising from multiple measurements per participant and are more flexible than repeated-measures Analysis of Variance [52]. LMMs deal with pseudoreplication and account for the fact that participants display individual differences in overall reaction time and accuracy.

In the detection task, RT increased significantly with log batch as evidenced by the positive regression slope (LMM, *β* = 224.705, *z* = 15.837, *p <* 0.001). Analogously, there was significant increase in ’real’ responses with log batch (*β* = 6.184, *z* = 9.343, *p <* 0.001) and a significant decrease in correct responses (*β* = −6.195, *z* = − 9.405, *p <* 0.001). We then used Wilcoxon Signed-Rank tests to compare RTs and performance for batch 60000 vs real MRIs across participants. No difference was found for RTs (*w* = 146, *p* = 0.672) and correct responses (*w* = 119, *p* = 0.252) but the proportion ’real’ was significantly higher for real MRIs than for generated ones (*w* = 0, *p <* 0.0001).

In the subjective rating task, RT increased significantly with log batch (*β* = 279.875, *z* = 11.377, *p <* 0.001). For the analysis of the ratings, a linear mixed model approach would not be not sufficient since the target variable was ordinal and the ’distances’ between neighbouring categories were unknown [53]. The Mean Opinion Scores displayed in Figure 4e were used for illustrative purposes. In line with the recommendations in [53] we used a multi-level proportional odds model with a logit link function which simultaneously deals with mixed effects and ordinal responses. To this end, we fit a Cumulative Link Mixed Model with Laplace approximation [54], wherein rate was used as the ordinal target variable, log batch as fixed effect and participant as a random effect with random intercept. We found a significant positive slope for log batch (*β* = 1.234, *z* = 17.94, *p <* 0.001) signifying higher ratings for later batches. The same statistical approach was applied to compare ratings for batch 60000 vs real MRIs, showing significantly higher ratings for real MRIs as compared to generated ones (*β* = 2.133, *z* = 7.748, *p <* 0.0001).

### 3.4. Image quality metrics and Deep QA model

Figure 5 shows the metrics calculated for the detection task images. An analogous analysis applied to the rating task images is depicted in Figure S3 (Supplementary Material). Figure 6b shows the corresponding result for the Deep QA model. Linear regression analyses of log batch on the metric were conducted to investigate whether the metric increases/decreases with batch number.

**Figure 5:**
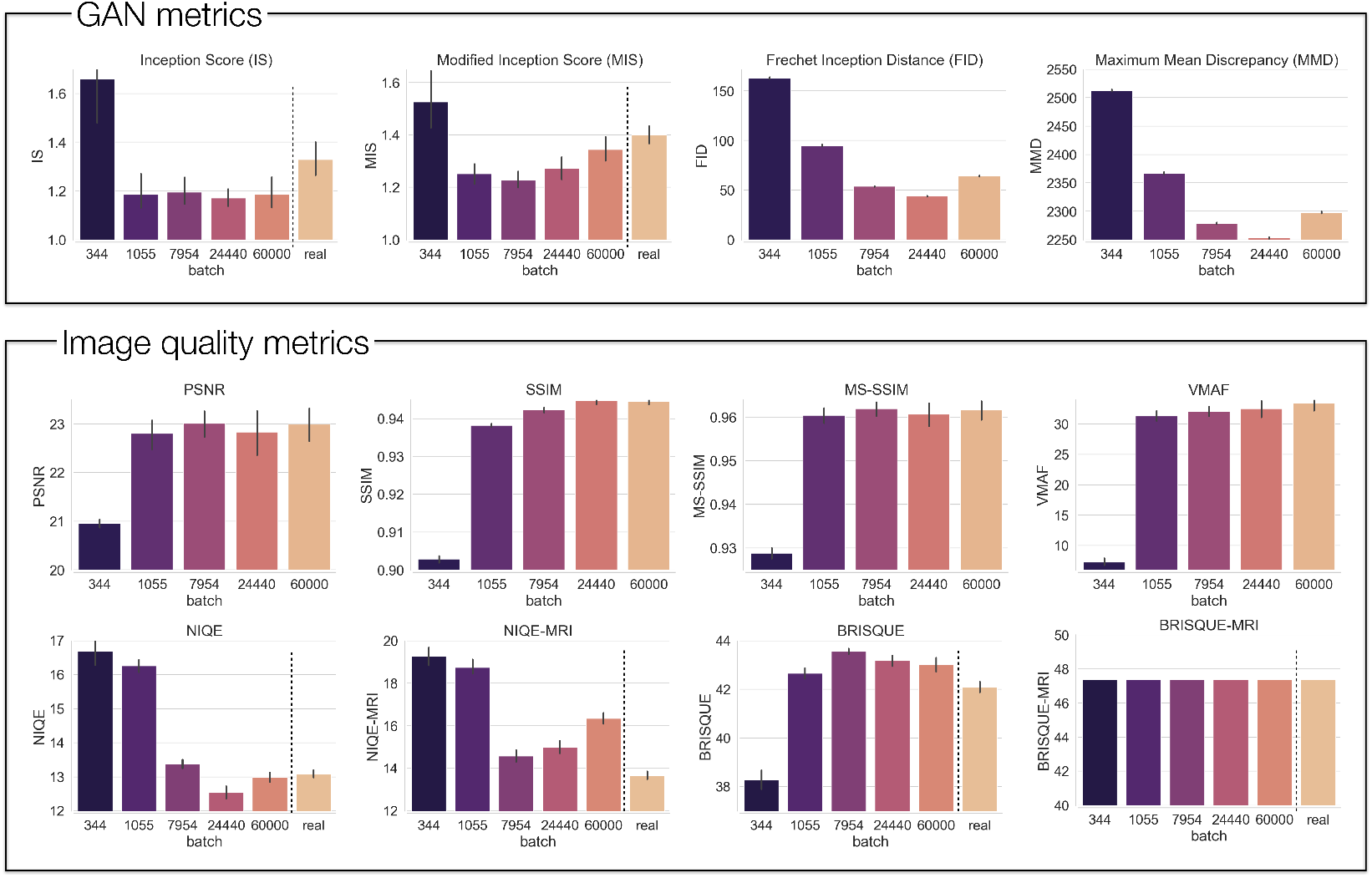
Image quality metrics applied to the 240 images used in the detection experiment. Different GAN and image/video metrics are shown, applied to generated images from different batches. If the metric does not require a reference, the value for real images is shown as well.

**Figure 6:**
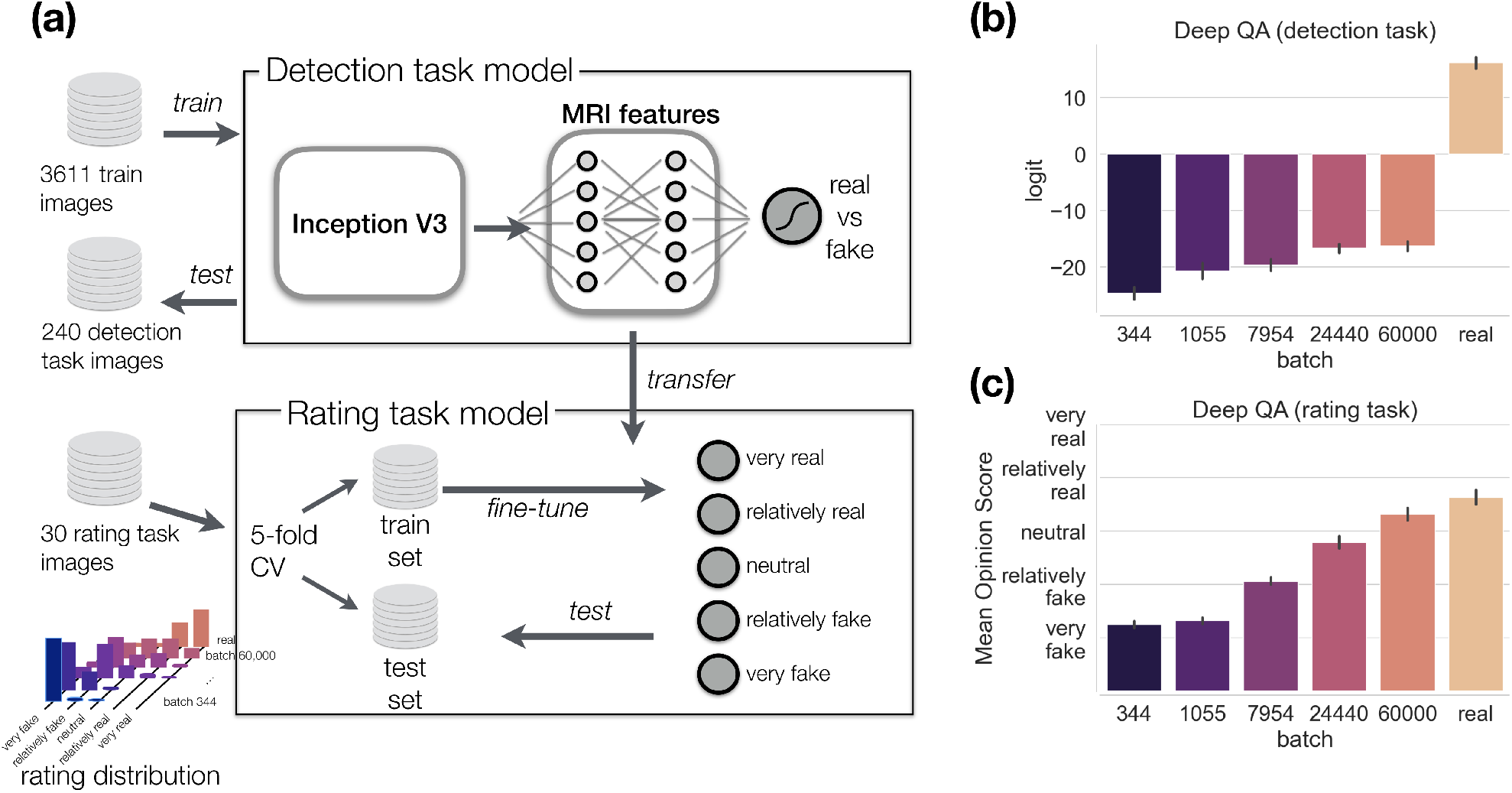
Deep QA model. (a) Model architecture. A pretrained InceptionV3 model was used followed by two dense layers (MRI features) trained on independent data. It was then fine-tuned on the rating data using cross-validation. (b) Logits of the predicted probabilities on the detection task images. (c) Mean Opinion Scores on the rating task images. Means and confidence intervals were calculated across 100 runs.

For the detection data, all regression slopes were significant (all p-values *<* 0.0001 except for MIS *p* = 0.0009) except for BRISQUE-MRI which gave a constant output and was discarded from further analysis. IS and MIS both decreased with batch number. Since they are quality metrics, the opposite pattern was expected. FID and MMD both decreased with batch number but images at batch 24400 were predicted to have a better quality than images at batch 60000 (Wilcoxon Rank Sum, FID *z* = − 12.217, *p <* 0.001; MMD *z* = − 12.217, *p <* 0.001), conflicting with the behavioral data. PSNR, SSIM, MS-SSIM, and VMAF increased with batch number. Of these, only VMAF increased monotonically, in line with the behavioral data, and SSIM increased monotonically except for the last batch. However, in neither case was the difference between batches 24400 and 60000 significant (Wilcoxon Rank Sum test, SSIM *p* = 0.493; VMAF *p* = 0.282). Although NIQE(-MRI) and BRISQUE(-MRI) showed overall regression trends in line with behavioral data, the pattern for batches 7954, 24400 and 60000 conflicted with human data. NIQE predicted a lower quality for the 60000 batch than for 24400 (*z* = − 2.611, *p* = 0.009). Except for VMAF, the Deep QA model was the only model for which predictions on the detection data changed monotonically with batch number, although the difference between batches 24400 and 60000 was not significant (*p* = 0.51).

We repeated the same analysis on the rating task data (Figure S3 and Figure 6b). Except for IS (*p* = 0.4424), MIS (*p* = 0.1943), and BRISQUE-MRI (*p* = 1.0), all regression slopes were significant (*p <* 0.001 except PSNR *p* = 0.0023). Again, VMAF increased monotonically and SSIM increased monotonically except for the last batch. However, in neither case was the difference between batches 24400 and 60000 significant (Wilcoxon Rank Sum test, SSIM p = 0.4647, VMAF p = 0.347). Deep QA was the only metric that showed a significant difference between these two batches (*z* = − 2.193, *p* = 0.0282).

To statistically compare the image quality metrics to the behavioral data, we performed Spearman’s rank correlation analyses for both tasks, shown in Table 1. For the detection task, we used the proportion ’real’ responses for each of the 240 images, averaged across participants, and correlated them with the metrics obtained on the same data (Figure 5 and Figure 6b). FID and MMD could not be subjected to correlation analysis because they estimate the distance between distributions rather than individual images. Since some metrics did not provide separate predictions for real images, we conducted two correlation analyses, one analysis that covered only generated images and another one that included real images (denoted as ’include real’ in the table). For the rating task, we used the Mean Opinion Scores for each of the 30 images, averaged across participants, and correlated them with the metrics obtained on the rating task data (Figure 6c and Figure S3).

**Table 1:** Correlation between behavioral data and image quality metrics reported as Spearman’s *ρ* and corresponding p-value, for the two behavioral tasks separately. Each analysis was performed twice, one time using only the data on the generated images, and a second time including the real data, too. For each analysis, the three metrics with the largest absolute correlation are highlighted in bold.

| Task |  | IS | MIS | PSNR | SSIM | MS-SSIM | VMAF | NIQE | NIQE-MRI | BRISQUE | BRISQUE-MRI | DeepQA |
| --- | --- | --- | --- | --- | --- | --- | --- | --- | --- | --- | --- | --- |
| Detection | $\rho$ | -0.250 | -0.016 | 0.421 | **0.754** | 0.459 | 0.404 | **-0.706** | **-0.536** | 0.465 | NaN | 0.516 |
|  | p-value | 0.001 | 0.845 | 0.000 | **0.000** | 0.000 | 0.000 | **0.000** | **0.000** | 0.000 | NaN | 0.000 |
| Detection (include real) | $\rho$ | 0.019 | 0.241 | - | - | - | - | **-0.519** | **-0.761** | -0.013 | NaN | **0.831** |
|  | p-value | 0.767 | 0.000 | - | - | - | - | **0.000** | **0.000** | 0.840 | NaN | **0.000** |
| Rating | $\rho$ | -0.206 | 0.112 | 0.705 | **0.925** | 0.657 | 0.504 | **-0.767** | -0.575 | 0.521 | -0.067 | **0.879** |
|  | p-value | 0.323 | 0.593 | 0.000 | **0.000** | 0.000 | 0.010 | **0.000** | 0.003 | 0.008 | 0.751 | **0.000** |
| Rating (include real) | $\rho$ | -0.065 | 0.104 | - | - | - | - | **-0.614** | **-0.722** | 0.294 | -0.052 | **0.912** |
|  | p-value | 0.733 | 0.585 | - | - | - | - | **0.000** | **0.000** | 0.114 | 0.786 | **0.000** |

Except for IS, MIS, BRISQUE, and BRISQUE-MRI, all metrics showed moderate to high correlation with the behavioral data. For the detection task, SSIM (*ρ* = 0.754), NIQE (*ρ* = −0.706), and NIQE-MRI (*ρ* = −0.536) were the three best performing metrics when the real data was excluded. When real images were included, the best performing metrics were Deep QA (*ρ* = 0.831), NIQE-MRI (*ρ* = −0.761) and NIQE (*ρ* = −0.519). Despite the fact that the Deep QA model performed the classification task better than humans (100% accuracy), it assigned higher probabilities to later batches which led to a ranking consistent with humans (Figure 6b). However, when real images were removed from the correlation analysis, the correlation for the Deep QA model plummeted to *ρ* = 0.516, indicating that the effect was largely driven by the difference between real and generated images.

For the rating task, SSIM (*ρ* = 0.925), Deep QA (*ρ* = − 0.879), and NIQE (*ρ* = − 0.767) were the best performing metrics when the real data was excluded. When real images were included, the best performing metrics were Deep QA (*ρ* = 0.912), NIQE-MRI (*ρ* = −0.722) and NIQE (*ρ* = −0.614).

## 4. Discussion

Human assessment is the current gold standard for judging image quality of images generated by GANs [8, 22]. However, the utility of human assessment is limited by economical factors, limited scalability of human raters to large datasets, and the limited sensitivity of the human eye to subtle statistical relationships across dozens or hundreds of images. By collecting both behavioral data, performing group analysis and spatial ICA, and surveying a whole range of image quality metrics, our goal was to identify a more automated approach for the assessment of generated images. These points will be expanded on in the next two subsections, followed by a broader discussion of our findings, the limits of GANs for image generation, and the ideal image quality metric.

### 4.1. Biological plausibility: group analysis and spatial ICA

Prior to performing spatial ICA, we investigated the degree of correspondence between real and generated MRIs at a voxel level. The correlation between generated and real images was 0.80 (Figure 3a), which was very similar to the correlation of 0.81 of real images with themselves (Figure 3b). However, the correlation of generated images with each other was 0.92. This suggests that while generated MRIs adequately approximate real MRIs, the latter have a larger amount of diversity, in line with a previous research [8]. One explanation for this is that the GAN lacks *capacity*, that is, it is simply unable to replicate the diversity found in real data because the data does not lie within the range of possible outputs of the generator. Alternatively, the GAN might simply have failed to learn the generative distribution of the MRIs, a possibility discussed in Section 4.3.

We then performed group analysis to investigate whether the GAN reproduces group differences. This is not a trivial question since the GAN is only trained to produce images that ’look like brains’. It is not trained to reproduce statistical regularities and the discriminator that guides the training process does not have access to all images simultaneously to facilitate such a learning. Analysis of group differences considered training separate GANs to characterize chronological age (young vs old) and sex (male vs female). The groups of generated images were then subjected to group analysis. We found that the age effect is fairly well reproduced (Figure 3c) while the replication of the gender effect was rather low (Figure 3d). The low reproduction of the gender effect might be due to the fact that gender differences contribute less variance to the grey matter signal than aging which involves global atrophy.

Furthermore, we performed spatial ICA to investigate whether the GAN reproduces structural brain networks known to support cognitive function. Again, the model is not explicitly trained to perform this. Performing spatial ICA on real and generated images separately and comparing the results we found a large degree of correspondence between the spatial maps (Figure 3e). We found spatial patterns reminiscent of the networks identified in other studies of ageing with source-based morphometry [55, 31]. This correlation between the spatial maps was high irrespective of the exact choice of the number of ICs.

### 4.2. Image quality metrics as a proxy for human assessment

To identify whether image quality metrics correspond well with human assessment, we collected human data in two different tasks (detection and rating) and calculated a series of image quality metrics for the same images. For the human data, we found a significant increase in ’real’ responses with batch number. Since better performance was associated with lower RT, this could not be explained in terms of a speed-accuracy trade-off. A similar pattern of results was found in the subjective rating task, with Mean Opinion Scores (MOS) increasing significantly with batch number and approaching the ratings for real images. As hypothesized, these results confirmed that changes in objective image quality operationalized by batch number are perceptually relevant and measurable. Behavioral measures converged towards the responses obtained for real images, but there remained a gap, indicating that our model was not consistently able to fool human experts.

To relate behavioral data to the image quality metrics, we calculated Spearman’s rank correlation between quality metrics and the proportion of ‘real’ responses in the behavioral data. The standard GAN metrics IS and MIS showed poor correspondence with human assessment, in line with earlier research [22], and only few metrics agreed well with behavior. Both NIQE (detection task *ρ* = − 0.706; rating task *ρ* = − 0.767) and SSIM (detection *ρ* = 0.754; rating *ρ* = 0.925) showed a good agreement. However, SSIM did not show a significant difference for batches 24400 and 60000, whereas NIQE even predicted the 60000 batch to be of lower quality. Our Deep QA models showed a moderate correlation with human data in the detection task (*ρ* = 0.516, rising to *ρ* = 0.831 when including real images) and a high correlation in the rating task (*ρ* = 0.879 and *ρ* = 0.912 when including real images). It is worth noting that only the rating model was explicitly trained on human responses, whereas the detection model was only trained to discriminate between real and generated images. Crucially, the Deep QA rating model was the only metric that showed a significant increase in quality ratings from the 24400 to the 60000 batch.

An interesting conclusion of this analysis is that most image quality metrics correlate with human behavior reasonably well for lower batch numbers (batches 344, 1055, 7954) but are less sensitive to image differences later in training (batches 24400 vs 60000). We believe this discrepancy might be due to most metrics being sensitive to *distortion* rather than *perceptual quality*, two different dimensions of image quality that have been explored theoretically in image restoration tasks [56]. *Distortion* refers to the deviation of a distorted image from a noise-free reference whereas *perceptual quality* measures the more elusive ’naturalness’ of an image regardless of a reference. In our experiment, informal evidence in line with this is a participant stating^2^ that they focused on image artifacts such as checkerboard patterns for lowquality images (i.e., distortion), but that it was rather subtle differences in luminance across the image would that made higher-quality generated images look fake (i.e., perceptual quality).

If human assessment of image quality was indeed governed by distortion and artifacts for earlier batches and a more intuitive understanding of what an MRI should look like for later batches, it is reasonable to believe that most of the metrics were sensitive to artifacts, hence explaining their good performance for early batches and bad performance for the last batch. The only model that consistently agreed with human data across all quality levels was the Deep QA model. The model was directly trained on the images and human responses which gave it the opportunity to learn the more subtle features that humans use for more high-quality generated images.

### 4.3. The limits of GANs

The promise of GANs is somewhat uncanny: After being exposed to a few hundred images it supposedly learns the brain image manifold in image space. A key question therefore is whether the GAN indeed learns (a) the generative distribution of MRIs, whether it learns (b) a different distribution, or (c) simply memorizes and reproduces the training data. Regarding (c), it turns out that memorization of data requires a large number of different samples [57]. Other indicators contra memorization are the fact that interpolation between noise vectors can produce novel and meaningful image variations [58] and the clear disparity seen between real and generated images [59].

In accordance with (a), GANs can closely approximate the underlying generative distribution for sufficiently large datasets [6]. Generalization beyond the training data, indicated by the emergence of novel combinations of features, has been shown in simple datasets [60]. However, there is doubt that the same holds for limited sample sizes [59]. For instance, the *generative support* (crudely speaking, the number of individually different images the GAN can produce) found for faces strongly depends on the model architecture, with estimates ranging from 160,000 to over a million faces. This casts doubt on option (a). Further support for (b) is given by the phenomenon of mode collapse, i.e. the tendency of GANs to concentrate its probability mass to a few modes. [61] showed that the relative frequency of attributes such as hair style for faces or room type for indoor scenes is distorted from the real proportion in the data. Furthermore, classifier performance deteriorates when trained on GAN images rather than real data, and [61] estimated that the effective dataset size of the GAN is 100x smaller than the training data.

The debate on the generative support of GANs is ongoing, but results so far favour alternative (b), that is, GANs probably approximate the true distribution rather than just memorizing the data, but the support of the GAN distribution is limited. This limitation can probably be partially counteracted by using a large and diverse training set and a sufficiently capable CNN architecture. For limited size datasets, data augmentation could be a viable tool for increasing set size and diversity.

### 4.4. The ‘ideal’ quality metric

Although the focus of this paper was to assess how well existing metrics agree with human data, it seems expedient to list other favourable properties that an ideal MRI quality metric should possess:

1. *Model agnostic*. The metric should be independent of the specific model architecture in order to enable quantitative comparison across models. This precludes e.g. the discriminator’s loss function to act as a metric.
2. *Reference free*. It should determine the quality of an image per se, without resorting to a reference image. This would make it applicable to both noise-to-image and image-to-image applications and also in situations wherein the ground truth is not available.
3. *Individual image ratings*. A measure of image quality should be provided for individual images rather than the whole dataset.
4. *Multi-dimensional*. Although MRIs are often depicted as a series of 2D slices, the slices together actually form a 3D image and an accurate assessment should take into account all spatial dimensions.
5. *Human performance*. The metric has to reproduce the perceptual assessment of expert viewers. Furthermore, there is evidence that image manipulations that are imperceptible to humans can have a significant effect on CNN predictions [62]. Therefore, we believe that human assessment should serve as a *lower sensitivity bound* when evaluating image metrics. In other words, if humans are sensitive to the difference between two types of images, so should be the metric – but the metric could be sensitive to differences that humans are not.

None of the surveyed metrics meets all of these criteria. All of the surveyed metrics are model agnostic but only IS, MIS, NIQE, BRISQUE, and the Deep QA model are reference free. Individual image ratings are not available for FID and MMD because they operate on the level of distributions rather than individual images. Although all metrics were applied to 2D image slices, IS, MIS, MMD and MMD can be seamlessly extended to 3D by deriving a feature embedding from a 3D CNN. Furthermore, PSNR can directly operate on 3D images and for VMAF the third spatial dimension can be encoded as a temporal dimension, although this constitutes a ’hack’ of the metric. The Deep QA model can be trained on 3D images using 3D convolutions. A difficulty in extending metrics to 3D images is the collection of experimental data. A bespoke experimental protocol for assessing 3D images wherein participants can e.g. rotate, zoom into and slice 3D MRI images would need to be developed.

### 4.5. Limitations

The usage of a specific model (noise-to-image GAN) with a specific type of data (GM maps) leads to the question whether our results are relevant to other model architectures (e.g. VAEs), hyperparameter choices (e.g. number of convolutional layers), or imaging modalities. We believe they are, since all CNN-based models use the same computational building blocks such as upscaling operations and convolutional layers that produce well-known artifacts known as checkerboard patterns [63, 49]. With regard to imaging modalities, we believe that our results for GM maps are relevant to other types of images such as T1 and T2-weighted. For instance, [64] developed a multi-modal CNN that performs liver segmentation in both MRI and CT images, and [65] reported successful transfer learning from structural MRIs to Diffusion Tensor Imaging (DTI) data. Both studies suggest that there different imaging modalities share visual commonalities.

Furthermore, behavioral results were collected using an online experiment rather than a controlled study in a lab environment. Participants performed the experiments in different environments using different display devices (e.g. laptop screen vs monitor). This potentially reduces the internal validity of our results, although it may increase their external validity. We believe that this limitation mostly affects group analyses. For instance, the seniority of a participant may be correlated with the display device they use, thereby confounding group differences between junior and senior experts. However, we performed only within-subject analyses. If anything, the fact that our findings are consistent across a variety of environments and display devices strengthens rather than weakens their validity. For instance, using Amazon Mechanical Turk, [66] replicated various visual effects such as Stroop effect and attentional blink that have been studied extensively in laboratory settings. Another limitation is that we only considered a noise-to-image model, no image-to-image model. In the latter, reference images are available which might improve the quality of metrics that rely on reference images such as SSIM.

### 4.6. Conclusion

As a practical recommendation for evaluating brain images produced by GANs, we propose that researchers assess both local properties (visual quality of individual images) and global properties (statistical regularities across the dataset) of the generated data.

With regard to global properties, the generated dataset should reproduce relevant group differences such as young vs elderly or male vs female, with the specific choice of relevant groups depending on the nature of the dataset and the research question. Sensitivity of the GAN to group differences can be investigated by either re-training it on separate groups, as done in this paper, or using a conditional GAN with the demographics as separate input variables. Furthermore, a realistic GAN should be able to reproduce the large-scale structural networks exposed by multivariate analyses across the dataset. We performed spatial ICA to verify that the GAN reproduces components such as the Default Mode Network. Other approaches such as Principal Component Analysis (PCA) or non-negative matrix factorization [67] could be used for the same purpose.

With regard to local properties, the generated brain image should ’look like’ a real one. This can be verified using both distortion and perceptual metrics, two dimensions of image quality [56]. As a distortion metric, SSIM stood out as a versatile, widely used image quality metric that does not require tuning to the MRI data [26]. It was sensitive to a variety of image distortions and can be therefore be expected to perform similarly in other types of imaging datasets. It is also widely available on different platforms (e.g. ssim in Matlab and TensorFlow). A similar performance was found for NIQE (niqe in Matlab). Either of them or both metrics could be reported as distortion measures. As a perceptual metric, our Deep QA rating model was the only metric that reproduced human data for higher quality images. Unfortunately, the model required human data to be trained. This defeats its purpose to some extent since we set out to escape the need to collect human data. Although speculative, a possible way out of this predicament is to pretrain a Deep QA model on a large, diverse dataset that integrates multiple data modalities [65, 64]. Ideally, this model could be used out-of-the-box for any new dataset. Optionally, it could be fine-tuned to the statistics of a new dataset without requiring human data.

Concluding, we propose a combination of local and global analyses for assessing the quality of generated images. For the time being, human assessment remains the gold standard for assessing individual images. In the future, we believe that a Deep QA model that has been trained to mimick human perceptual assessments can pave the way for quick and cost-effective image quality assessments that will accelerate GAN research and improve its validation.

## Acknowledgements

We acknowledge the support of the Supercomputing Wales project, which is part-funded by the European Regional Development Fund (ERDF) via Welsh Government. KAT was supported by the Guarantors of Brain (G101149).

## Author contributions

MT conceptualized the study, collected human data and performed the statistical analyses. RC trained the GAN. MT and KT wrote the manuscript. All authors reviewed and approved the manuscript.

## Footnotes

1 http://mialab.mrn.org/software/gift

2 the participant disclosed their participation in the experiment to the author in a personal email

## References

[1] R. A. Poldrack, K. J. Gorgolewski, Making big data open: data sharing in neuroimaging, Nature Neuroscience 17 (2014) 1510–1517.

[2] H. Peng, W. Gong, C. F. Beckmann, A. Vedaldi, S. M. Smith, Accurate brain age prediction with lightweight deep neural networks, Medical Image Analysis 68 (2021) 101871.

[3] M. Sajjad, S. Khan, K. Muhammad, W. Wu, A. Ullah, S. W. Baik, Multi-grade brain tumor classification using deep CNN with extensive data augmentation, Journal of Computational Science 30 (2019) 174–182.

[4] G. Mohan, M. M. Subashini, MRI based medical image analysis: Survey on brain tumor grade classification, Biomedical Signal Processing and Control 39 (2018) 139–161.

[5] V. Sorin, Y. Barash, E. Konen, E. Klang, Creating Artificial Images for Radiology Applications Using Generative Adversarial Networks (GANs) – A Systematic Review, Academic Radiology 27 (2020) 1175–1185.

[6] I. J. Goodfellow, J. Pouget-Abadie, M. Mirza, B. Xu, D. Warde-Farley, S. Ozair, A. Courville, Y. Bengio, Generative Adversarial Nets, in: Z. G. Weinberger, M. Welling, C. Cortes, N. D. Lawrence, K. Q. (Eds.), Advances in Neural Information Processing Systems 27, Curran Associates, Inc., 2014, pp. 2672–2680.

[7] D. P. Kingma, M. Welling, Auto-encoding variational bayes, in: 2nd International Conference on Learning Representations, ICLR 2014 - Conference Track Proceedings, International Conference on Learning Representations, ICLR, 2014.

[8] A. U. Hirte, M. Platscher, T. Joyce, J. J. Heit, E. Tranvinh, C. Federau, Diffusion-Weighted Magnetic Resonance Brain Images Generation with Generative Adversarial Networks and Variational Autoencoders: A Comparison Study, arXiv (2020) 2006.13944.

[9] C. K. Chong, E. T. W. Ho, Synthesis of 3D MRI Brain Images with Shape and Texture Generative Adversarial Deep Neural Networks, IEEE Access 9 (2021) 64747–64760.

[10] S. U. H. Dar, M. Yurt, L. Karacan, A. Erdem, E. Erdem, T. Ç ukur, Image Synthesis in Multi-Contrast MRI with Conditional Generative Adversarial Networks, IEEE Transactions on Medical Imaging 38 (2019) 2375–2388.

[11] D. Nie, R. Trullo, J. Lian, C. Petitjean, S. Ruan, Q. Wang, D. Shen, Medical Image Synthesis with Context-Aware Generative Adversarial Networks, in: Medical Image Computing and Computer Assisted Intervention - MICCAI 2017. MICCAI 2017. Lecture Notes in Computer Science, Springer, Cham, 2017, pp. 417–425.

[12] D. Abramian, A. Eklund, Generating fMRI volumes from T1-weighted volumes using 3D CycleGAN, arXiv abs/1907.0 (2019).

[13] X. Gu, H. Knutsson, M. Nilsson, A. Eklund, Generating Diffusion MRI Scalar Maps from T1 Weighted Images Using Generative Adversarial Networks, Lecture Notes in Computer Science 11482 (2019) 489–498.

[14] Y. Chen, F. Shi, A. G. Christodoulou, Y. Xie, Z. Zhou, D. Li, Efficient and Accurate MRI Super-Resolution Using a Generative Adversarial Network and 3D Multi-level Densely Connected Network, in: Medical Image Computing and Computer Assisted Intervention – MICCAI 2018, Springer, Cham, 2018, pp. 91–99.

[15] T. M. Quan, T. Nguyen-Duc, W.-K. Jeong, Compressed Sensing MRI Reconstruction Using a Generative Adversarial Network With a Cyclic Loss, IEEE Transactions on Medical Imaging 37 (2018) 1488–1497.

[16] X. Yi, E. Walia, P. Babyn, Generative adversarial network in medical imaging: A review, arXiv e-prints 1809.07294 (2018).

[17] C. Bermudez, A. J. Plassard, T. L. Davis, A. T. Newton, S. M. Resnick, B. A. Landman, Learning Implicit Brain MRI Manifolds with Deep Learning, Proceedings of SPIE–the International Society for Optical Engineering 10574 (2018).

[18] K. Kazuhiro, R. A. Werner, F. Toriumi, M. S. Javadi, M. G. Pomper, L. B. Solnes, F. Verde, T. Higuchi, S. P. Rowe, Generative Adversarial Networks for the Creation of Realistic Artificial Brain Magnetic Resonance Images, Tomography (Ann Arbor, Mich.) 4 (2018) 159–163.

[19] A. Volokitin, E. Erdil, N. Karani, K. C. Tezcan, X. Chen, L. Van Gool, E. Konukoglu, Modelling the Distribution of 3D Brain MRI Using a 2D Slice VAE, in: Lecture Notes in Computer Science, volume 12267 LNCS, Springer Science and Business Media Deutschland GmbH, 2020, pp. 657–666.

[20] G. Kwon, C. Han, D.-s. Kim, Generation of 3D Brain MRI Using Auto-Encoding Generative Adversarial Networks, in: MICCAI 2019: Medical Image Computing and Computer Assisted Intervention, pp. 118–126.

[21] C. Han, L. Rundo, K. Murao, T. Noguchi, Y. Shimahara, Z. A. Milacski, S. Koshino, E. Sala, H. Nakayama, S. Satoh, MADGAN: unsupervised medical anomaly detection GAN using multiple adjacent brain MRI slice reconstruction, BMC Bioinformatics 2021 22:2 22 (2021) 1–20.

[22] F. Calimeri, A. Marzullo, C. Stamile, G. Terracina, Biomedical Data Augmentation Using Generative Adversarial Neural Networks, in: P. Verschure, A. Villa, A. Lintas, S. Rovetta (Eds.), Artificial Neural Networks and Machine Learning - ICANN 2017. ICANN 2017. Lecture Notes in Computer Science, Springer, Cham, 2017, pp. 626–634.

[23] C. Han, L. Rundo, K. Murao, Z. A. Milacski, K. Umemoto, H. Nakayama, S. Satoh, GAN-based Multiple Adjacent Brain MRI Slice Reconstruction for Unsupervised Alzheimer’s Disease Diagnosis, in: In International Meeting on Computational Intelligence Methods for Bioinformatics and Biostatistics. Lecture Notes in Computer Science, Springer, Cham, 2019, pp. 44–54.

[24] K. H. Kim, W. Do, S. Park, Improving resolution of MR images with an adversarial network incorporating images with different contrast, Medical Physics 45 (2018) 3120–3131.

[25] D. Yang, B. Liu, L. Axel, D. Metaxas, 3D LV Probabilistic Segmentation in Cardiac MRI Using Generative Adversarial Network, in: Statistical Atlases and Computational Models of the Heart. Atrial Segmentation and LV Quantification Challenges, Springer, Cham, 2019, pp. 181–190.

[26] Z. Wang, A. C. Bovik, H. R. Sheikh, E. P. Simoncelli, Image quality assessment: From error visibility to structural similarity, IEEE Transactions on Image Processing 13 (2004) 600–612.

[27] N. Luo, J. Sui, A. Abrol, J. Chen, J. A. Turner, E. Damaraju, Z. Fu, L. Fan, D. Lin, C. Zhuo, Y. Xu, D. C. Glahn, A. L. Rodrigue, M. T. Banich, G. D. Pearlson, V. D. Calhoun, Structural brain networks match intrinsic functional networks and vary across domains: a study from 15000+ individuals, Cerebral Cortex 30 (2020) 5460–5470.

[28] M. Shafto, L. K. Tyler, M. Dixon, J. R. Taylor, J. B. Rowe, R. Cusack, A. J. Calder, W. D. Marslen-Wilson, J. Duncan, T. Dalgleish, R. N. Henson, C. Brayne, F. E. Matthews, The Cambridge Centre for Ageing and Neuroscience (Cam-CAN) study protocol: a cross-sectional, lifespan, multidisciplinary examination of healthy cognitive ageing, BMC neurology 14 (2014) 204.

[29] J. R. Taylor, N. Williams, R. Cusack, T. Auer, M. A. Shafto, M. Dixon, L. K. Tyler, Cam-CAN, R. N. Henson, The Cambridge Centre for Ageing and Neuroscience (Cam-CAN) data repository: Structural and functional MRI, MEG, and cognitive data from a cross-sectional adult lifespan sample, NeuroImage 144 (2017) 262–269.

[30] J. Ashburner, A fast diffeomorphic image registration algorithm, NeuroImage 38 (2007) 95–113.

[31] K. A. Tsvetanov, R. N. A. Henson, P. S. Jones, H. Mutsaerts, D. Fuhrmann, L. K. Tyler, J. B. Rowe, The effects of age on resting-state BOLD signal variability is explained by cardiovascular and cerebrovascular factors, Psychophysiology 58 (2021) e13714.

[32] M. Arjovsky, S. Chintala, L. Bottou, Wasserstein Generative Adversarial Networks, in: Proceedings of the 34th International Conference on Machine Learning, pp. 214–223.

[33] I. Gulrajani, F. Ahmed, M. Arjovsky, V. Dumoulin, A. C. Courville, Improved Training of Wasserstein GANs, in: Proceedings of the 31st International Conference on Neural Information Processing Systems, Curran Associates Inc., Long Beach, California, USA, 2017, p. 5769–5779.

[34] G. Stoet, PsyToolkit: A software package for programming psychological experiments using Linux, Behavior Research Methods 42 (2010) 1096–1104.

[35] G. Stoet, PsyToolkit: A Novel Web-Based Method for Running Online Questionnaires and Reaction-Time Experiments, Teaching of Psychology 44 (2017) 24–31.

[36] R. Whelan, Effective analysis of reaction time data, Psychological Record 58 (2008) 475–482.

[37] M. Frid-Adar, I. Diamant, E. Klang, M. Amitai, J. Goldberger, H. Greenspan, GAN-based synthetic medical image augmentation for increased CNN performance in liver lesion classification, Neurocomputing 321 (2018) 321–331.

[38] L. Xu, K. M. Groth, G. Pearlson, D. J. Schretlen, V. D. Calhoun, Source-based morphometry: The use of independent component analysis to identify gray matter differences with application to schizophrenia, Human Brain Mapping 30 (2009) 711–724.

[39] A. Borji, Pros and cons of GAN evaluation measures, Computer Vision and Image Understanding 179 (2019) 41–65.

[40] C. Lee, S. Woo, S. Baek, J. Han, J. Chae, J. Rim, Comparison of objective quality models for adaptive bit-streaming services, in: 2017 8th International Conference on Information, Intelligence, Systems & Applications (IISA), volume 2017, Institute of Electrical and Electronics Engineers Inc., 2017, pp. 1–4.

[41] J. Jia Deng, W. Wei Dong, R. Socher, L.-J. Li-Jia Li, K. Kai Li, L. Li Fei-Fei, ImageNet: A large-scale hierarchical image database, in: 2009 IEEE Conference on Computer Vision and Pattern Recognition, IEEE, 2009, pp. 248–255.

[42] S. Barratt, R. Sharma, A Note on the Inception Score, arXiv (2018) 1801.01973.

[43] S. Gurumurthy, R. K. Sarvadevabhatla, V. B. Radhakrishnan, DeLi-GAN : Generative Adversarial Networks for Diverse and Limited Data, Proceedings - 30th IEEE Conference on Computer Vision and Pattern Recognition, CVPR 2017 2017-Janua (2017) 4941–4949.

[44] M. Heusel, H. Ramsauer, T. Unterthiner, B. Nessler, S. Hochreiter, GANs Trained by a Two Time-Scale Update Rule Converge to a Local Nash Equilibrium, Advances in Neural Information Processing Systems 2017-Decem (2017) 6627–6638.

[45] A. Gretton, K. M. Borgwardt, M. J. Rasch, B. Schölkopf, A. Smola, A Kernel Two-Sample Test, Journal of Machine Learning Research 13 (2012) 723–773.

[46] Z. Wang, E. P. Simoncelli, A. C. Bovik, Multi-scale structural similarity for image quality assessment, in: Conference Record of the Asilomar Conference on Signals, Systems and Computers, volume 2, pp. 1398–1402.

[47] Z. Li, A. Aaron, I. Katsavounidis, A. Moorthy, M. Manohara, Toward A Practical Perceptual Video Quality Metric, Netflix Technology Blog (2016) Jun 6.

[48] A. Mittal, A. Krishna Moorthy, A. Conrad Bovik, No-Reference Image Quality Assessment in the Spatial Domain, IEEE Transactions on Image Processing 21 (2012) 4695–4708.

[49] G. Yang, Y. Cao, X. Xing, M. Wei, Perceptual Loss Based Super-Resolution Reconstruction from Single Magnetic Resonance Imaging, in: Lecture Notes in Computer Science, volume 11632,Springer Verlag, 2019, pp. 411–424.

[50] R. Zhang, P. Isola, A. A. Efros, E. Shechtman, O. Wang, The Un-reasonable Effectiveness of Deep Features as a Perceptual Metric, in: Proceedings of the IEEE Conference on Computer Vision and Pattern Recognition (CVPR), pp. 586–595.

[51] C. Rorden, M. Brett, Stereotaxic display of brain lesions, Behavioural Neurology 12 (2000) 191–200.

[52] D. A. Magezi, Linear mixed-effects models for within-participant psychology experiments: an introductory tutorial and free, graphical user interface (LMMgui), Frontiers in Psychology 6 (2015) 2.

[53] C. Keeble, P. D. Baxter, A. J. Gislason-Lee, L. A. Treadgold, A. G. Davies, Methods for the analysis of ordinal response data inmedical image quality assessment, British Journal of Radiology 89 (2016).

[54] R. H. B. Christensen, ordinal—Regression Models for Ordinal Data, 2019.

[55] K. Liu, S. Yao, K. Chen, J. Zhang, L. Yao, K. Li, Z. Jin, X. Guo, Structural Brain Network Changes across the Adult Lifespan, Frontiers in Aging Neuroscience 9 (2017) 275.

[56] Y. Blau, T. Michaeli, The Perception-Distortion Tradeoff, in: Proceedings of the IEEE Conference on Computer Vision and Pattern Recognition (CVPR), pp. 6228–6237.

[57] V. Nagarajan, C. Raffel, G. Brain, I. J. Goodfellow Google Brain, Theoretical Insights into Memorization in GANs, in: 32nd Conference on Neural Information Processing Systems.

[58] A. Radford, L. Metz, S. Chintala, Unsupervised representation learning with deep convolutional generative adversarial networks, in: 4th International Conference on Learning Representations, ICLR 2016 - Conference Track Proceedings, International Conference on Learning Representations, ICLR, 2016.

[59] S. Arora, Y. Zhang, Do GANs actually learn the distribution? An empirical study, arXiv (2017) 1706.08224.

[60] S. Zhao, H. Ren, A. Yuan, J. Song, N. Goodman, S. Ermon, Bias and Generalization in Deep Generative Models: An Empirical Study, Advances in Neural Information Processing Systems 2018-Decem (2018) 10792–10801.

[61] S. Santurkar, L. Schmidt, A. Madry, A Classification-Based Perspective on GAN Distributions, in: International Conference on Learning Representations (ICLR).

[62] J. Zhang, C. Li, Adversarial Examples: Opportunities and Challenges, IEEE Transactions on Neural Networks and Learning Systems 31 (2020) 2578–2593.

[63] Y. Sugawara, S. Shiota, H. Kiya, Checkerboard artifacts free convolutional neural networks, APSIPA Transactions on Signal and Information Processing 8 (2019) 1–9.

[64] K. Wang, A. Mamidipalli, T. Retson, N. Bahrami, K. Hasenstab, K. Blansit, E. Bass, T. Delgado, G. Cunha, M. S. Middleton, R. Loomba, B. A. Neuschwander-Tetri, C. B. Sirlin, A. Hsiao, Automated CT and MRI Liver Segmentation and Biometry Using a Generalized Convolutional Neural Network, Radiology: Artificial Intelligence 1 (2019) 180022.

[65] K. Aderghal, A. Khvostikov, A. Krylov, J. Benois-Pineau, K. Afdel, G. Catheline, Classification of Alzheimer Disease on Imaging Modalities with Deep CNNs Using Cross-Modal Transfer Learning, in: Proceedings - IEEE Symposium on Computer-Based Medical Systems, volume 2018-June, Institute of Electrical and Electronics Engineers Inc., 2018, pp. 345–350.

[66] M. J. C. Crump, J. V. McDonnell, T. M. Gureckis, Evaluating Amazon’s Mechanical Turk as a Tool for Experimental Behavioral Research, PLoS ONE 8 (2013) e57410.

[67] A. Anderson, P. K. Douglas, W. T. Kerr, V. S. Haynes, A. L. Yuille, J. Xie, Y. N. Wu, J. A. Brown, M. S. Cohen, Non-negative matrix factorization of multimodal MRI, fMRI and phenotypic data reveals differential changes in default mode subnetworks in ADHD, NeuroImage 102 (2014) 207–219.

